# Visualizing micro-nanoplastics in the human brain: Early evidence for roles in microvascular pathology

**DOI:** 10.64898/2026.09.14.751595

**Authors:** Elaine L. Bearer, Marcus A. Garcia, Laurissa Barela, Andres Collazo, Cathleen F. Martinez, Taylor W. Uselman, Gary A. Rosenberg

## Abstract

Our environment has become progressively contaminated with non-biological materials, especially synthetic carbon polymers, plastics. Unsurprisingly some of these make it into the human body as detected by chemical analyses, yet it is unknown whether they cause harm or are innocent bystanders. Building on our observations of glossy deposits and unusual non-biological fluorescent particles in blood vessels, we aimed to determine whether these objects represented plastics. To do this we prepared plastics-enriched pellets from brain and subjected them to pyrolysis gas-chromatography/mass spectroscopy (py-GC/MS), electron microscopy and confocal laser scanning microscopy. Py-GC/MS confirmed the presence of 10 different plastics. Contents of pellets were examined by thin-section EM. We then used laser scanning confocal microscopy to acquire hyperspectral profiles of each type of plastic in suspensions. We found fluorescent particles in all pellets from brains. All 12 plastics in the calibration standard fluoresced. We obtained hyperspectral profiles of single species plastics from industry: Polyethylene, polypropylene and polystyrene. Controls included chemical analysis of brain storage buffer, which had no detectable plastics; and imaging water only and areas on slides lacking tissue. Abundant particles with emission profiles similar to polyethylene and polypropylene decorated the walls of both arterioles and venules. These particles were coincident with glossy deposits we first observed in white matter, suggesting that both features represent plastics. In summary our results demonstrate that synthetic polymers exhibit detectable fluorescence, and particles with similar spectral properties are visible in histologic sections. This opens the door to investigating correlations between plastics and pathology.

**Highlights:**

- In brain histology, deposits that do not stain for hemosiderin were common adjacent to blood vessels.
- Pyrolysis gas-chromatography/mass-spectroscopy identified 10 different plastics in brain.
- Plastics in calibration standards fluoresced when excited by 405nm laser light.
- Synthetic plastics and particles in brain fluoresced with similar unique hyperspectral profiles.
- Particles that fluoresce like plastics were coincident with glossy deposits seen by brightfield.

## Introduction

Increasing levels of contamination of the environment from micro/nanoplastics are a growing concern for life in our modern era^1,2^. Recent reports attest to their presence within the human body, including testis^3^, placenta^4^ and even the brain^5,6^. In an effort to determine whether these plastics associate with histopathology, we set out to develop an optical technique that could detect them in microscopic examination of tissue sections. This is how most tissue pathology is diagnosed. Such an approach would allow us to detect tiny nanoparticles in the context of tissue architecture at cellular scale.

Pyrolysis gas chromatography/mass spectroscopy (py-GC/MS) is emerging as a powerful chemical technique for plastics detection and identification^7–9^, yet it requires a large amount of tissue (500mg) and does not preserve histopathological context. Other techniques, such as Raman spectroscopy or MassSpec-based microscopy can detect molecular signatures of specific plastics in tissue sections^10,11^, but the resolution is typically at a 10µm level, these techniques are destructive and little information about the pathology of surrounding tissue is obtained. In contrast, fluorescence microscopy can achieve detection of bright particles below the wavelength of visible light (500nm), and even approach nanoscale resolution, with pixel sizes less than 100nm and even smaller, especially when particles are bright enough to apply a point-spread function (PSF)^12^. Yet whether plastics are naturally fluorescent or can be detected over tissue background or other fluorescent tissue components is unknown.

Unique excitation/emission profiles of fluorescence from different plastic types could serve to differentiate them in tissue, just as charge/mass of their fragments differentiates plastic types by mass spectroscopy. By obtaining the excitation/emission profiles of synthetic plastics we can discover if plastics are naturally fluorescent and compare pure plastic subtypes with particles in enriched pellets of digested tissue and in histologic sections of brains that were the original source for the plastics-enriched pellet. In this way, we hoped to identify fluorescent particles as plastics within the specimens, and to explore the possibility that spectral profiles might even determine the type of plastic the fluorescing particles represent.

We reported that post-mortem brains from cases dying with cognitive impairment with high plastics burdens in brain also displayed many unusual tiny fluorescent particles in blood vessels of the choroid plexus, as detected by laser scanning confocal microscopy^13,14^. These particles had similar, but unique, excitation/emission profiles which differed from those of any other known fluorescent species in brain. We then found that similarly fluorescent particles resided in vessel walls of subcortical arterioles where substantial pathological changes were evident^15^. However, whether these fluorescent particles were plastics was not clear. Although the literature suggests that plastics fluoresce, their theoretical chemical formula predicts they would not^16^. With powerful lasers used for confocal imaging, the fluorescence of synthetic plastics and plastics isolated from tissue can be tested.

Our long history applying confocal microscopy to image fluorescent, 100nm diameter, polystyrene microspheres as probes for axonal transport^17–21^ provided the experience and technical skills needed to explore the fluorescent properties of these tissue-resident particles in detail. We also previously developed high-resolution fluorescence microscopy to study actin dynamics in cells and *in vitro*, with the ability to detect and record activity of 6-8nm fluorescently labeled filaments^22,23^, demonstrating that objects in the size range of nanoplastics found in tissues (predicted to be 40x200nm^5,6^) can be observed with optical techniques. Here we apply this knowledge and these skills to determine if the fluorescent particles we reported in human brain are plastics. This study uses multiple orthogonal techniques to determine whether the fluorescent particles observed exhibit spectral properties similar to that used as the polymer standard.

## Methods

### Post-mortem human brains

Post-mortem brain tissue was obtained from two consented participants enrolled in the University of New Mexico on-going NIH-supported MarkVCID and Alzheimer’s Disease Research Center (ADRC) programs, including one individual with a clinical Diagnosis of Binswanger’s disease (Case 1) and a second with neuropathologically confirmed Alzheimer’s disease (Case 2). UNM IRB determined that post-mortem tissue is not human subject research. Consents for brain removal, retention and use for research were approved by the Office of Medical Investigator (OMI) according to New Mexico State Law, and further reviewed by the UNM-HSC Human Tissue Review Board. Post-mortem consent was obtained from the participant’s legally authorized representative after deaths, according to New Mexico state law.

### Neurodegenerative disease workup

Neuropathology examination was performed for diagnostic purposes and the residual brain tissue deposited in the UNM Brain Bank for storage and sharing through the New Mexico Alzheimer’s Research Center. Following autopsy, brains are fixed and maintained in 20% formalin for 3-5 weeks until gross examination by a neuropathologist. Blocks harvested for histopathological diagnosis were subsequently sectioned and stained for this project. Tissue blocks were submitted for dehydration and paraffin embedding followed by sectioning, mounting on glass slides and histochemical staining at TriCore Reference Laboratories, or for sectioning and specialized immunohistochemistry (IHC) staining in UNM’s Human Tissue Repository, both CLIA-approved diagnostic labs, according to their standard clinically-proven protocols, including standard hematoxylin and eosin (H&E) ^24^. Additional histochemical stains were according to standard histopathology including: Perl’s Prussian Blue (for hemosiderin) with potassium ferrocyanide and nuclear fast red counterstain ^24^, and immunohistochemistry (IHC) for matrix-metalloproteinases, MMP1 (Primary, Rb polyclonal, Proteintech 10371-2-AP at 1:400) and MMP 10 (Primary, Rb polyclonal Abcam ab38930 at 1:200). Diagnostic stains for Alzheimer’s disease markers are routinely performed at TriCore. For IHC of MMPs, After heating and deparaffinization, antigen retrieval and endogenous peroxidase block, sections on slides were stained on the Ventana Discovery Platform with: Detection System - HRP Multimer: Anti-HQ HRP (Ventana 760-4820) is applied for 12 minutes; Chromogen: DISCOVERY Purple Kit (Ventana 253-4857) is applied for 8 minutes; Counterstain: Hematoxylin (Ventana 760-2021) is applied for 4 minutes. Post-Counterstain/Bluing: Bluing solution (Ventana 760-2037) is applied for 4 minutes). Histochemistry was

### Plastics Enriched Pellet

Brain tissues (∼500mg) dissected from regions adjacent to those submitted for neuropathologic examination were collected for microplastic analysis. White matter was dissected from cortical tissue and submitted separately from gray matter. Tissue was weighted and photographs taken after dissection. To assess potential contamination from the storage fluid, 1mL aliquots of the formalin storage solution were collected, evaporated in the pyrolysis cup, processed in parallel to the tissue, and then subjected in series by pyrolysis gas-chromatography/mass-spectroscopy (py-GC/MS) analysis (Agilent 8890 GC/5975 MS system (Agilent, Santa Clara, CA) with an EGA/PY-3030D Pyrolysis Unit (Frontier Labs, Koriyama, Japan). Approximately 500 mg from each specimen was selected for processing, sliced or minced using stainless steel instruments to increase surface area, and then placed in a 5mL glass vial. Room contamination was minimized by performing all steps in a laminar flow hood and avoiding plastics by glass containers. Three times the tissue volume (for 500 mg, this is 1.5mL) of 10% aqueous potassium hydroxide (KOH) was added. All samples were incubated at 40°C for 3-5 days, vortexed once a day for 2 minutes to increase mixing and digestion efficiency. After incubation, the contents of the vials were removed and individually transferred to 1.8mL polycarbonate ultracentrifuge tube with 200 µL of 100% ethanol added to promote particle separation. Samples were then vortexed until appearing homogeneous. Samples were centrifuged at 100,000xg for 4 hours at 4 °C. Following ultracentrifugation, the supernatant was removed and stored, and the pellet was washed with 500µL of 100% ethanol three times by resuspending with vortexing and re-pelleting at 10,000x g for 5 min. After the last wash the pellet was allowed to dry with the tube loosely covered for 24h at room temperature in a laminar flow hood to minimize room air contamination. After drying, the pellet was weighed and transferred to a glass vial for storage or other studies as previously described^3–6^.

Pellets from the tissue were subdivided for parallel analysis by 1) pyrolysis gas chromatography-mass spectrometry (py-GC/MS); 2) electron microscopy; and 3) hyperspectral confocal laser scanning microscopy. True procedural blanks for py-GC/MS, as described ^3–6^ , which included storage fluid and true blanks, were treated with KOH in polycarbonate ultracentrifuge tubes and subjected to the same ultracentrifugation process to see if anything could be collected from them. The evaporated residue was then subjected to py-GC/MS in sequence with the tissue pellet.

### Pyrolysis Gas Chromatography-Mass Spectrometry (py-GC/MS)

Plastic-enriched pellets were analyzed by py-GC/MS in duplicate at the University of New Mexico, using established protocols previously applied to the analysis of microplastics in human placenta, testis, and brain tissues^3–6^. Samples were run in sequence with certified microplastics calibration standards MPs-CaCO_3_ from Frontiers Lab (Frontier Laboratories, Fukushima, Japan). Calibration standards containing 12 polymers including polyethylene (PE, polypropylene (PP), polystyrene (PS), polyvinyl chloride (PVC, polycarbonate (PC), poly(methyl methacrylate) (PMMA), polyethylene terephthalate (PET), acronitrile-butadiene-styrene (ABS), styrene-butadiene rubber (SBR), polyurethane (PU), nylon 6 (N6), and nylon 66 (N66) were weighed out into different pyrolysis cups at 5 different weights (0.1mg, 0.2mg, 0.5mg, 2mg, and 4mg) together with storage buffer and procedural blanks run sequentially from an autosampler. The plastic-enriched pellets were weighed and approximately 1mg of each dried pellet was weighed into a stainless steel pyrolysis Eco-Cup (Frontier Laboratories, Fukushima, Japan). Samples were then analyzed using a Frontier EGA/PY-3030D multi-shot pyrolyzer set to “single shot” mode, directly coupled to an Agilent 8890 GC/5977 MS system (Agilent Technologies, Santa Clara, CA). A UAMP Column kit (Frontier Laboratories, Fukishima, Japan) designed for microplastic analysis was added to the GC/MS and was calibrated and configured for the analysis, set up in accordance with the Agilent manufacturer’s guidelines and integrated into the instrument’s control software. The instrument parameters included the Inlet Temperature (300° C), MS Source (230° C), starting oven temperature (40° C), MS Quad (150° C), Aux-2 Temperature (280° C), Column-1 Flow Cal Pressure (1.0), Foreline Pressure (<80), Inlet pressure (14.3 mL/min), and Column Flow (1 mL/min). Polymer identification was performed using F-Search MPs Software (Frontier Laboratories, Fukishima, Japan) using the MS patterns from the GC peaks automatically shunted into the MS and charge/mass (m/z) of multiple fragments within specific peaks compared to known standards for each type of plastic using the Frontier software, F-search (Frontier Laboratories, Fukishima, Japan).

### Electron microscopy

To examine confirm our ability to detect small plastic particles by confocal microscopy, we performed negative stain EM on commercially available 50nm polystyrene (PS) beads. A 1:100 dilution of the bead suspension (50 nm Nanoparticles, at 1% solid content (Phosphorex, Hopkinton, MA) in double-distilled water (DDW). The suspension was triturated and droplets (10µL) were placed on a freshly exposed, naïve paraffin sheet for 1 minute, followed by 2x 10µL droplets of DDW and a final 10µL droplet of 1% aqueous uranyl acetate (Electron Microscopy Sciences, Morgantown, PA) for 1 minute, according to our previously published methods for analyzing transport vesicles or herpesvirus^25–28^. All procedures were performed under a cover. To monitor for contaminants in the DDW or the UA solution, or from the environment during processing, droplets with only DDW or with DDW followed by UA were processed in parallel to droplets containing PS beads side-by-side within the same laboratory environment. Formvar-carbon-coated copper 200 mesh grids (Electron Microscopy Sciences, EMSciences, Morgantown, PA) that had been subjected to glow-discharge in a Denton Vacuum DV-502 evaporator in the Marine Biological Laboratory (MBL) Central Microscopy Facility in Woods Hole, MA. A glow discharge attachment was utilized in this workflow to perform mild low-vacuum plasma treatments. Exposure to glow discharge modifies carbon-stabilized Formvar support films to make them hydrophilic, which is essential for ensuring that aquatic, marine, or macromolecular biological specimens adhere smoothly to the grid surface during preparation. After reaching -100 Torr negative pressure, the filament charge was turned up until it emitted a purple and this was sustained for 3-5min. Glow-discharged grids were placed on top of each specimen-containing droplet for 1-3 minutes, then moved successively through DDW droplets for ∼30 sec each, and finally placed on UA droplet for 1 min. The UA was sucked off each grid with a damp triangular piece of Whatman filter paper No. 1, dried under a heat lamp and placed on filter paper in a small glass petri dish labeled in lead pencil. Grids were imaged in a JEOL CX200 electron microscope in the Central Microscopy Facility. Some grids of PS were not stained with UA and imaged both by EM and the grid subsequently mounted on glass slides, covered with a No 1.5 Gold Seal glass coverslip, and imaged by confocal microscopy (see below).

To examine the contents of the plastic-enriched pellets, we performed thin-section electron microscopy. Four plastics-enriched pellets from gray and white matter dissected from 2 different brains were examined. Plastic-enriched pellets (∼1mg) were resuspended in double distilled water (DDW) (100 µL). 50 µL of each suspension was mixed with equal parts of warm 2% gelatin in DDW, centrifuged into a pellet in Eppendorf tubes, and was left to gel on ice. The pellets were cut out of the tubes and processed in microwave processor with secondary fixation with 2.5% glutaraldehyde in 200mM sodium cacodylate buffer to fix the gelatin, followed by ice cold aqueous 1% OsO4, dehydrated in increasing ethanol concentrations, and embedded in Spurr’s resin. No acetone or any solvent other than ethanol was used. Sections were cut with a Diamond knife and mounted on Formvar coated EM grids. The sections were stained with 1% aqueous uranyl acetate followed by lead citrate. Images were collected with a Zeiss EM 10C operating at 80 kVa.

### Plastic suspensions for fluorescence analysis

Synthetic polymers were obtained as follows: polyethylene (PE), average M_w_∼4,000 by GPC, average M_n_∼1,700 by GPC, average diameter 149µm, highly polydisperse, range 5-300 µm (Sigma Aldrich, St. Louis, MO), polypropylene (PP) 10 mesh with 0-20 µm diameter particles (Magerial Sciences, Sheridan, WY), polystyrene (PS) nanoparticles (50 nm diameter, sized) (Phosphorex, Hopkinton, MA), and calibration standards from standards MPs-CaCO_3_ from Frontiers Lab (Frontier Laboratories, Fukushima, Japan).

For confocal, synthetic polymers and particles in the plastic-enriched preparations were suspended in DDW at approximately 1 mg/100µL. Suspensions were dropped onto pre-washed clean glass slides (3 or 6 mL) and covered with a #1.5 thickness glass coverslips (0.17µm) as recommended for confocal microscopy because the high numerical aperture (NA) objectives are optically corrected specifically for this thickness. The edges of the No. 1.5 coverslips were sealed with clear nail polish. To monitor contamination from the water, environment or pre-existing on the glassware, multiple control preparations were prepared in parallel under the same conditions and imaged: slides alone, slides with coverslips, and slides with 6 µL droplets of DDW and coverslipped with identical confocal parameters as for plastic suspension or tissue sections. Slides were stored in a covered slide box After preparation and before imaging, slides and coverslips were cleaned again with Zeiss lens.

### Lambda scans and multiplexed images by confocal laser-scanning fluorescence

Suspensions and tissue sections were imaged by laser scanning confocal microscopy. Confocal images were collected on the inverted Zeiss LSM 980 microscope with Fast Airyscan 2 module and a 32-channel GaAsP detector array in the Biological Imaging Facility in the Beckman Institute at Caltech. This confocal has 7 laser lines (405, 440, 488, 514, 561, 594 & 639 nm wavelengths) and capability for white light imaging with a photomultiplier tube (PMT). Objectives included Zeiss Plan Apochromat 4x, 10x, 20x, dry 40x (N.A. 0.95), and 63x (N.A. 1.4) and 100x (N.A. 1.3) oil immersion. Once the region of interest was found at low magnification, immersion oil was applied and the objective switched to a higher magnification lens. The 405nm laser was used for lambda scanning, with emissions collected at ∼8.8nm step sizes from low 400 to 700nm wavelengths for 32 images. Using the Zoom function, a pixel size of <100nm could be achieved. Lambda scans were analyzed in Zen Blue using the Unmix function by encircling regions of interest and measuring the intensities across the series of emissions. A rectangular area with no signal was selected for background measurement. Typically this background had no measurable emission at 405nm excitation. The table of intensity values in arbitrary units was copied and saved as a csv file. Histograms of intensity versus wavelength were generated in Excel. PMT images were multiplexed using the multi-alkali T-PMT detector (detection wavelength 300-900nm). To capture images for size measurements, we identified the ideal sampling density by calculating the Nyquist rate for the 63x 1.4N.A. objective and refractile index of 1.5255, excitation 405 nm, emission 450 nm, with a pinhole of 1 Airy Unit, which gave 0.18^2^ micron (https://svi.nl/Nyquist-Calculator). We estimated that the average minimal size of a plastics particle isolated from brain would 40 x 200nm, as previously reported^6^ and that most would be clustered into larger aggregates due to their hydrophobicity. We therefore used a 3.1x Zoom to acquire 85 nm^2^ pixel sizes, which is less than half the larger dimension of the target particle. With a pixel size of 0.085 microns (a.k.a. 85 nm), our target size threshold of 200 nm converts to 2.35 linear pixels.

### Spectral Profiling of Fluorescent Particles and Tissue Features

To perform hyperspectral analysis from lambda scans, regions of interest (ROIs) were manually drawn in Zeiss’s ZEN software Unmix tab. For suspended plastic samples, ROIs were defined as circles or ellipsoids for suspected plastic particles and as rectangles for background non-particle containing regions. Regions were selected based on their appearance in a stacked image across all emissions. Size of the region of interest was determined by zooming up on the screen and selected a circle that included as little background as possible. The area of these regions ranged from 2-4 µm diameter. Pixels sizes ranged from 0.085-0.263µm. For 50nm polystyrene nanoparticles, a 20x objective with 3.2x Zoom (pixel size 0.129 µm) with pinhole set at 1 airy unit was sufficient to detect beads based on their luminosity when excited by the 405 laser at 4% power, detector Gain at 724V, 0 Offset and digital gain of 1.

For histological sections, ROIs were similarly drawn for plastic-like aggregates fluorescing in the 410-490nm range under 405 laser exciting, and for non-tissue background regions, with additional measurements from tissue features including collagen and blood identified by their typical structure visible in the PMT image or in images from longer wavelengths. Multiple ROIs were selected from plastic-like aggregates within a sample. Arbitrary intensities were obtained in Zen Blue unmix window, and the table of arbitrary intensity values copied and pasted into Google sheets, downloaded as a csv and opened in excel for calculations and graphing. Average fluorescence intensities were calculated for each ROI at each wavelength step in the emission spectrum and spectral intensity profiles inserted as line graphs in excel. For background subtracted intensity graphs, the background was measured in a large area devoid of fluorescent particles and similarly transported into the excel spreadsheet containing the ROI data. In some instances, background intensity for each ROI at each wavelength was subtracted from the particle intensity value. In Zeiss ZEN software, intensity in arbitrary units (a.u.) is relative to the internal raw digital pixel gray values recorded by the camera sensor or photomultiplier tube (PMT), and scaled by the specific settings for laser power, detector gain, and exposure time. Background intensity typically hovered within 0.1 a.u. of zero, whereas intensity of synthetic plastic particles and of those putative plastics isolated from tissue often exceeded 50 or even 200 a.u. in the same dataset image.

For some measurements, intensity profiles were adjusted in Excel using Min-Max normalization. For Min-Max normalization, each ROI’s intensity measurement was subtracted by the minimum intensity across ROIs in that image and then divided by the intensity range (maximum - minimum) of ROIs in that image, producing values between 0 and 1. For samples with multiple ROIs of similar type (e.g., multiple plastic-like aggregates or single industry plastic standards), intensities at each wavelength step were averaged across ROIs, and standard deviations calculated and shown on graphs as error bars. Individual ROI spectral profiles were plotted as line graphs for all samples; when multiple ROIs of the same feature type (i.e., plastic-like particles) were obtained, averaged profiles with error bars representing standard deviations were also plotted. For each graph, all ROIs were measured from the same multidimensional image containing 32 separate images across the emission range, with each image obtained by the detector for 8.8nm continuous nanometer wavelengths (ie, 419-427nm for channel 1). The photomultiplier tube (PMT) was used to collect transmitted white light for detection of all objects in the field as not all fluoresce.

### Particles within histologic sections of brain fluoresce with plastic-like profiles

Tissue sections with various stains were imaged both with lambda scans, as for plastic particles, as well as with 2- or 3-channel imaging using 3 lasers and the PMT. The selection of excitation/emission wavelengths were informed by the lambda scan results. Two- to four-color images were collected of the same field with: 1) 405nm Ex. (3.5%)/410-472nm Em.; 2) 488nm Ex. (0.2%)/516-543nm Em.; and 3) 561nm (0.2%)/578-667nm Em. With or without 4) a DIC image (PMT). Pinhole was maintained at 1 A.U. for each laser. Detector Gain and Offset were adjusted for each image based on “Best fit” for visualization in the Zen software. Typically, scan mode was unidirectional frame with 16 averages for a ∼2 sec pixel time and 20 sec frame time with scan speed set at 7. For some images these parameters were adjusted to improve image quality or magnification. To monitor for background fluorescence from the uninvolved tissue, or the mounting media, lambda scans were performed on regions lacking tissue, regions further distant from blood vessels, collagen fibers identified by their distinctive appearance in microscopy, neuromelanin granules identified by their unique shape, intracellular location and brown pigment in large adrenergic neurons of substantia nigra and locus coeruleus, and lipofuscin in large motor neurons in the pons.

## Results

### Histopathology of brain discovered non-biological objects previously unrecognized

Glossy, yellowish deposits were observed adjacent to blood vessels in subcortical white matter by histopathology of a person who died with Binswanger’s disease. White matter hyperintensities, characteristic but not diagnostic of this disease, were present in annual FLAIR MRIs for 7 years prior to death in this person’s brain. To assess the pathological basis for these WMHs, we selected subcortical regions that had displayed this abnormality in life to dissect and embed for histopathological investigation (**Fig. 1**). Sections from tissue blocks from these regions were stained for matrix-metalloproteinase 1 (MMP1) and 10 (MMP10), to address the hypothesis that these enzymes would contribute to loss of vascular integrity and edema, which likely produced the hyperintense MR signal. While histologically vessels appeared intact, abnormal glossy yellowish deposits were seen, particularly noticeable in regions of white matter ischemic change when stained with a new purple chromogen with hematoxylin counter stain^15^. The smooth muscle in the media of this vessel seemed normal, stained for MMP1, while perivascular macrophages, stained for MMP10, appeared more abundant that normal. The glossy deposits, visible in both these immunohistochemical stains, were many different sizes ranging from sub-micron to several microns in diameter.

**Figure 1.**
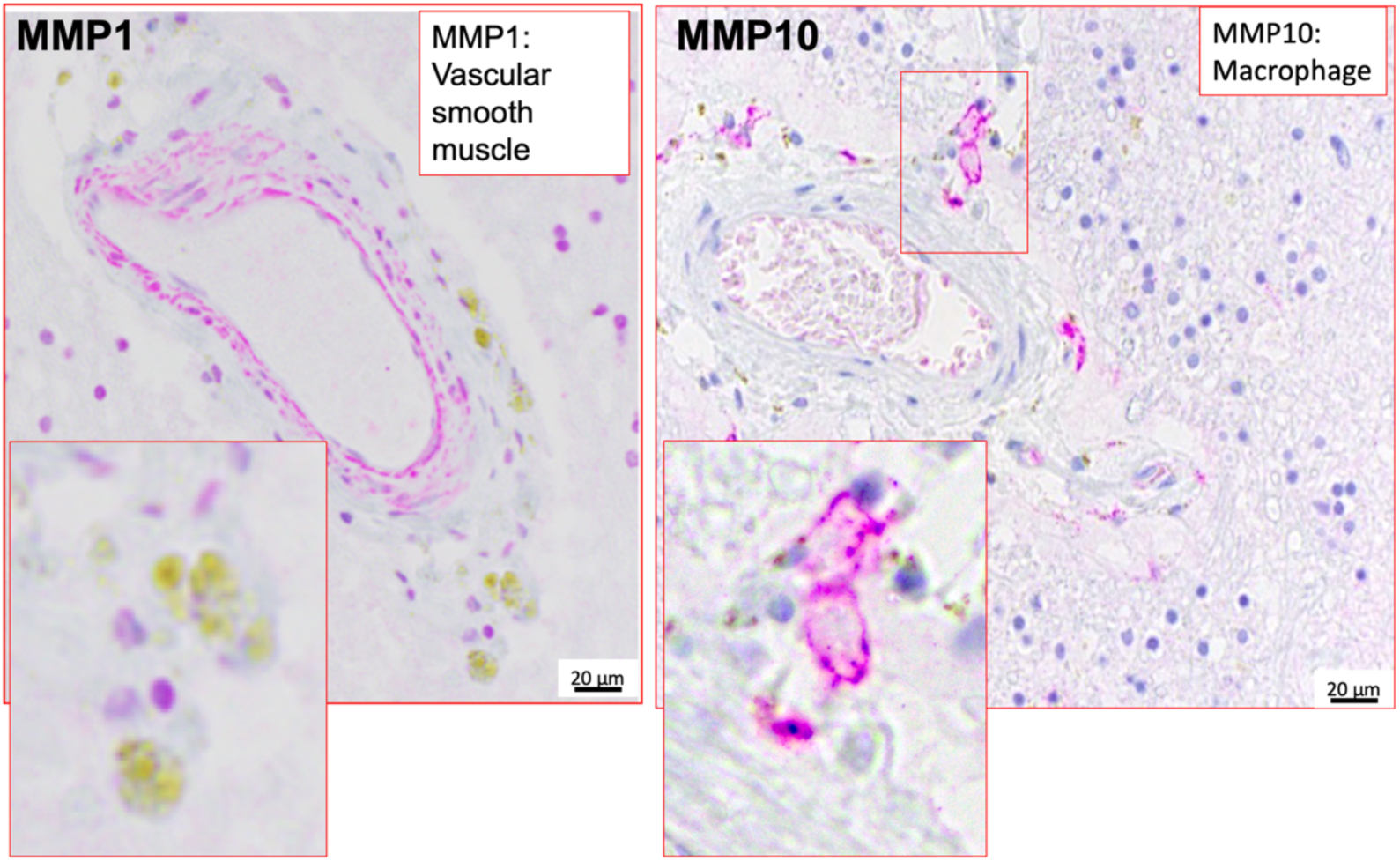
Brightfield images of yellowish glossy deposits within blood vessels walls of subcortical white matter. (**A**) Example image of a section stained by IHC for MMP1 with secondary purple chromogen. MMP1 stains the muscle layer, while unidentified yellow appears in particles clustered outside the vessel walls in perivascular connective tissue. Red square identifies one area shown enlarged in the inset. Mag bar = 20 µm. (**B**) Example image of a section stained by IHC with MMP10 primary antibody, secondary antibody labelled for purple chromogen. MMP10 stained perivascular macrophages. In this example, yellowish deposits are found outside the vessel and around the MMP10 positive cells, not within them. Squares indicate one of the positive areas selected for enlargement in the inset. Mag bar = 20µm.

The initial neuropathological assumption was that these deposits were residuals from extravasated RBCs subsequent to a distant micro-hemorrhage, where the hemoglobin had been degraded to hemosiderin over time. This idea could be tested by examining sections from the same block after staining for hemosiderin with Prussian blue coupled with nuclear fast red as counterstain^24^ (**Fig. 2**). As expected, hemosiderin in sections of liver, from a case with hemosiderosis that was mounted on the same slide as the brain sections, stained blue with Prussian blue. The glossy deposits did not, demonstrating they were not hemosiderin. Virtually all blood vessels in this representative section displayed yellow-brown glossy deposits that were not hemosiderin (**Fig 2 B-F**), showing that this effect was wide-spread throughout the region of brain that displayed WMH. These regions also had white matter ischemic changes, evidenced by foamy macrophages and loss of myelin (not shown). Similar results were obtained from multiple different brain regions in this and other cases.

**Figure 2.**
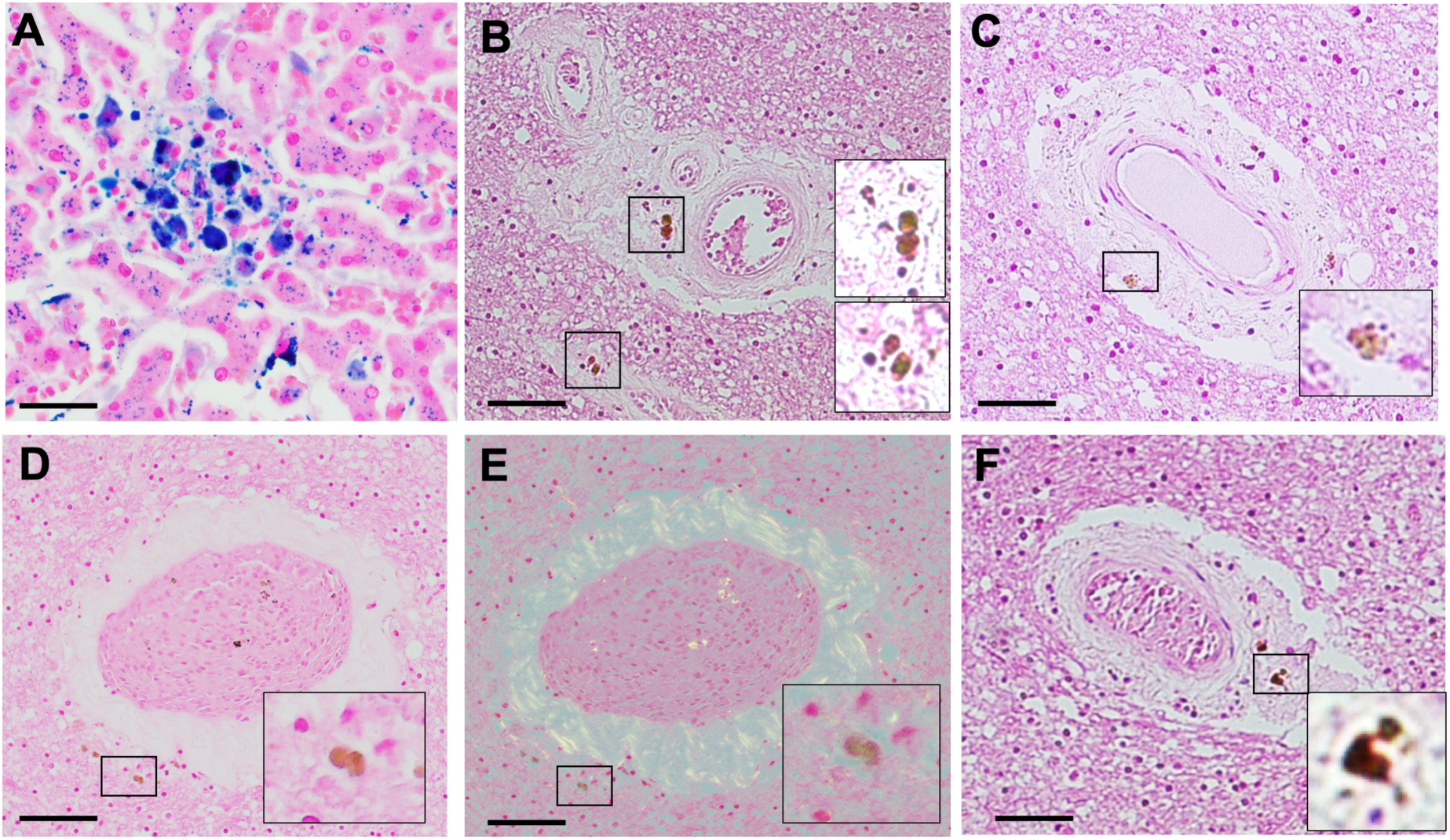
Brightfield images of Prussian-blue stained sections from subcortical white matter. (**A**) Control section of liver from a person with hemosiderosis, where hemosiderin collects in the liver (blue). Counter stain (pink) was nuclear fast red. The liver sections were places on the same slide as the brain sections and processed simultaneously. Mag bar = XX (**B-F**) Blood vessels in subcortical white matter displaying deposits adjacent to the outer vessel wall. Both arterioles with a smooth muscle layer (**B, C, F**) and venules (**E-F**, same vessel) lacking this layer display these particulate deposits which do not stain blue for hemosiderin. Polarization imaging emphasizes the collagen surrounding the venule (**F**). Deposits lie within the adventitia or slightly beyond it into the adjacent perivascular space or brain parenchyma. No blood vessels were found lacking deposits, although the numbers and location differed slightly.

### Chemical analysis to identify plastics in brain tissue

To ascertain if this case harbored measurable amounts of plastics and of which type, 500mg of adjacent white matter tissue was submitted for py-GC/MS according to previously published protocols ^3–6^ (**Fig. 3**). Our tissue blocks are typically 4 mm thick and 1.5 x 1cm square, approximately weighting 0.6gm, comparable to the 0.5gm chunk of brain dissected for plastics analysis. The tissue was first digested in 10% aqueous KOH expected to reverse formalin cross-linking, particulate matter collected by centrifugation, the pellet washed in 100% ethanol and 1mg aliquots processed in duplicate through the Agilent py-GC/MS as described for other brain material ^5,6^ (**Fig. 3A-C**). As a control for plastics possibly emanating from the storage container or from the lab environment during processing, we collected storage buffer and processed it in parallel to the tissue blocks. (**Fig. 3A & B**). To minimize contamination from containers or the environment, glass containers were used whenever possible and all procedures were undertaken in a laminar flow cell culture hood.

**Figure 3.**
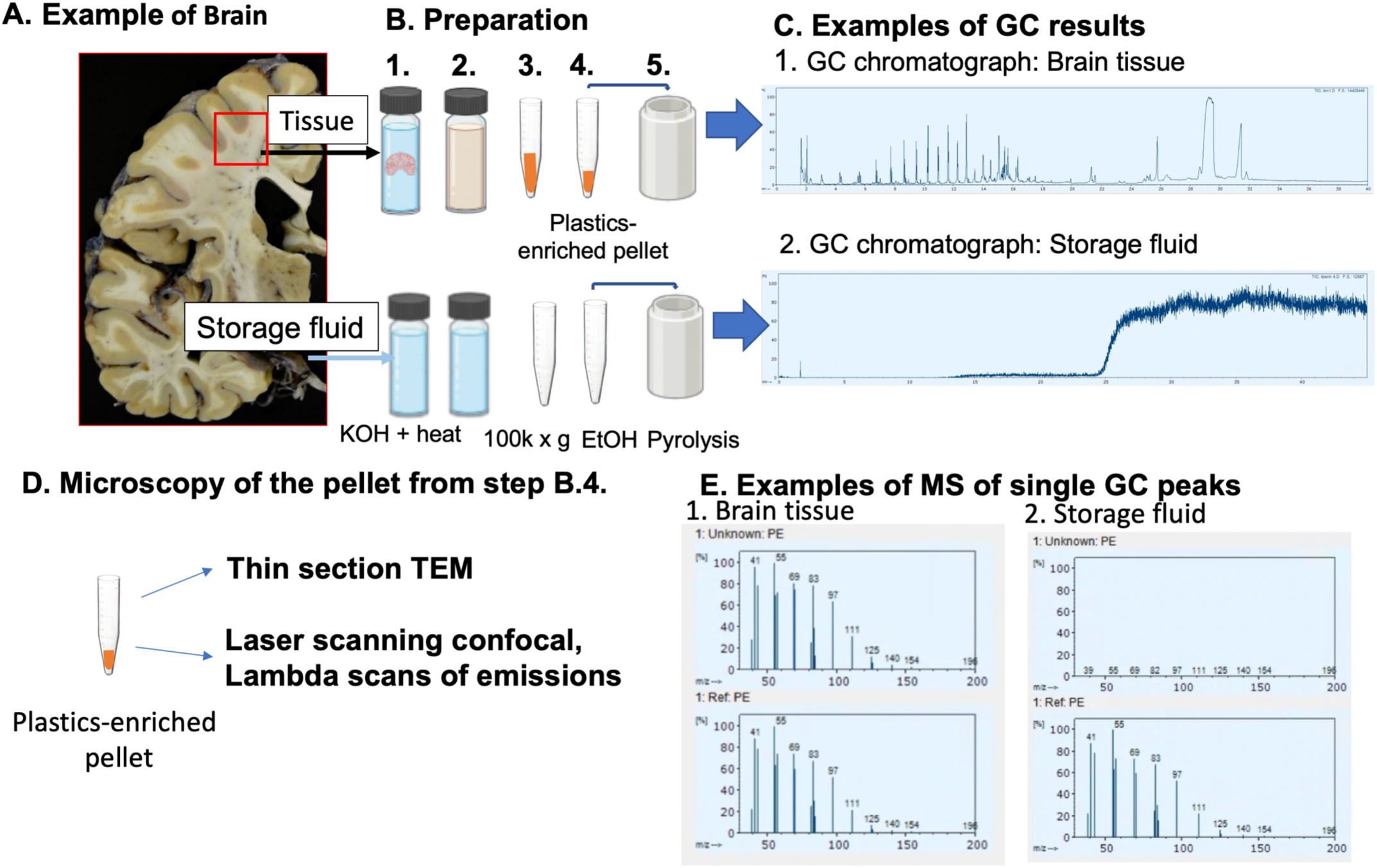
Diagram and results from plastics-enrich pellets of brain tissue. (A) An example of one of the gross brain slices used for this analysis showing the location of the tissue specimen (red square) and diagramming the storage fluid used as internal control for plastics contamination. (B) Illustration of the processing steps with 1. tissue digestion in KOH at 40° C heat for 72h; 2. Solublized material after digestion; 3. Pellet after 100k x g centrifugation that separated solid from soluble digested material and considered to contain plastics; 4. Washes of the pellet three times in ethanol (EtOH); 5. Pyrolysis in the metal pyrolysis cup. (C) Examples of the gas chromatographs for: 1. Brain tissue; and 2. Storage fluid. Note lack of peaks in the left half of the storage fluid chromatograph. These are the retention peaks in the brain gas chromatograph that, when analyzed by mass spectroscopy, display signatures of specific plastic types. (D) Cartoon of how the plastics-enriched pellet is subdivided to different analytical procedures. IN addition to py-GC/MS, material in other aliquots of the same pellet were also subjected to EM and confocal imaging. (E) Example of mass spectroscopy results from the GC peak with retention time consistent with polyethylene (PE) with: 1: Brain tissue, top, result from the tissue with unknown PE; bottom results from a reference PE sample run in sequence; 2. Storage fluid results, top, with no MS signal from the effluent with the same retention time as that sampled in 1; bottom, results from a reference PS sample run in sequence.

Typically 500mg of white matter yielded a post-digestion pellet of around 50-80mg, or 10-16 percent, whereas gray matter (10-37 mg). A relatively high pellet mass suggested the recovered material contained residual mass in addition to isolated plastic particles. The gas chromatogram for all brain tissue samples gave multiple retention time (RT) peaks (**Fig. 3C**), while only one small early peak appeared from the storage buffer sample which probably represented carbon dioxide from the initiation of gas flow when the py-GC/MS starts. RTs from this GC column run under these parameters are consistent, giving specific peaks for particular plastics. Thus each peak at a specific RT can be further analyzed by MS and compared to the peaks produced by known plastics in calibration standards, run in sequence to the duplicates of the brain samples with unknown plastics in them, or to archival data acquired under identical conditions. The remainder of the pellet was divided and either submitted for various types of EM examination or for confocal imaging (**Fig. 3D**).

To identify plastic types of plastics by py-GC/MS, calibration standards which contain a set of 12 plastics listed in the methods section, was run in sequence through the system. Individual peaks from the gas chromatogram were shunted into the mass spectroscope and subjected to fragmentation. The mass/charge ratio (m/z) of resultant fragments were measured by Mass Spectroscopy, which gives a particular profile for each specific plastic type. When residual tissue matrix is retained in the sample, this can alter the MS profile and interfere with quantification. Shown in **Figure 3E** are representative MS profiles from the unknown brain tissue of the 13.5 sec RT peak (**Fig. 3E1** top) and the corresponding peak from PE standards (**Fig.3E1,** bottom). Similarly, the same RT peak from the GC of the storage buffer is shown (**Fig. 3E**2, top), which has no signal, and the PE in the standards, with is the classical PE profile (**Fig. 3E**, bottom). This procedure detected at least 10 different plastics in the brain samples. Here we focused on polyethylene, polypropylene, and polystyrene for further fluorescence characterization.

### Thin section electron-microscopy of the plastics-enriched pellets

To monitor the contents of the plastics-enriched pellet used for chemical identity and for optical imaging, we embedded aliquots of each pellet for thin section electron-microscopy (**Fig. 4**). Plastics in the resuspended plastics-enriched pellet were centrifuged into warm gelatin to hold them in place for embedding. After cooling to solidify the gelatin, a block was cut out of the tube, trimmed for EM sectioning and dehydrated for plastic embedding. Blocks were fixed in gluteraldehyde, stained in Osmium tetroxide, dehydrated and embedded in Spurr’s. To avoid solubilizing the plastics during dehydration and possibly losing them, we used only ethanol instead of the usual solvents. Sections were stained with 1% aqueous uranyl acetate and lead citrate^26,27,29–31^.

**Figure 4.**
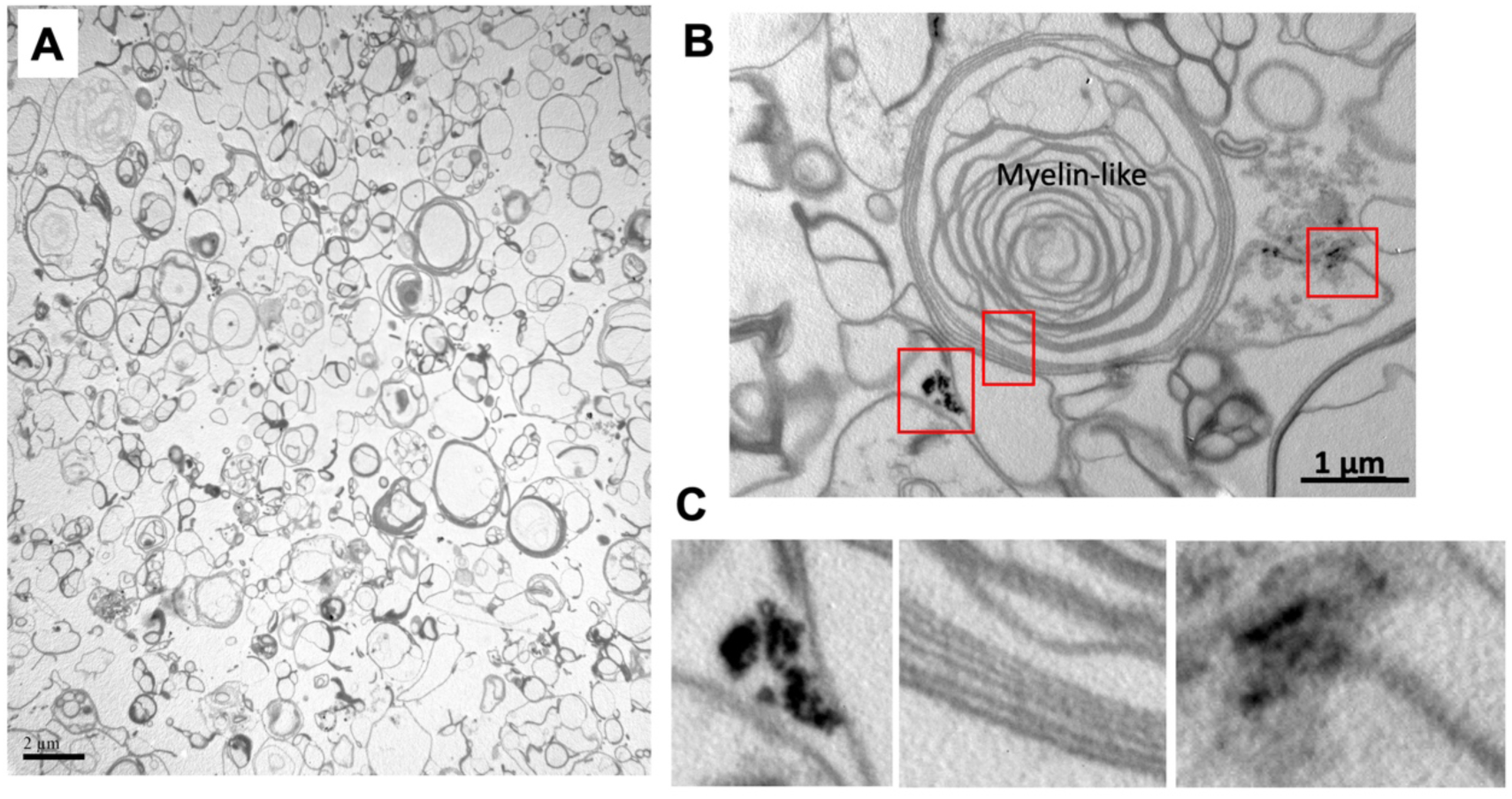
Representative thin section electron-microscopy of one of the plastic-enriched pellets. (A) Low magnification image of pellet contents. Note numerous round vesicles. Mag bar = 2 µm. (B) Higher magnification of the same sample in a different field. Multi-lamellar vesicles undigested resemble myelin. Red squares indicate regions that are arbitrarily enlarged to show osmicated particles that may represent plastics and the multilayered lipid membranes. Mag bar = 1µm.

Sections of blocks from 4 different plastics-enriched pellets demonstrated abundant lipid membranes (**Fig.4**). Some osmicated shards resembled the size and shape of plastic-like particles previously described by whole mount, unstained EM of ethanol and ultrasound treated plastics-enriched pellets from brain^5,6^. These shards are reported to range about 40nm by 200nm. Since plastics are well known to take up osmium, these black shards in the thin section may indeed represent the plastics found in the pellet by py-GC/MS, although they could also be osmium precipitates. The presence of abundant lipid membranes, some myelin-like, suggested that quantification of the py-GC/MS would be confounded and not reliable. However, plastics typing from the MS of specific, individual GC peaks by comparisons to known plastic types seemed to us to be valid.

### Hyperspectral analysis of particles from brain and comparison of lambda scans to that of plastics

To investigate whether the glossy deposits observed in the brains that contained plastics by py-GC/MS, and see if these were the same as the fluorescent particles we had reported previously^13–15^, we performed lambda scans across the visible wavelengths by laser scanning confocal microscopy on plastic particles from plastics-enriched pellets after suspension in double-distilled water. We used double distilled water as it is purified in glass, piped through metal and stored in glass bottles, whereas deionized water is filtered and stored in plastic containers. Suspensions were mounted on pre-washed glass microscope slides and covered with clean glass coverslips and sealed for laser confocal scanning.

We performed lambda scans on the suspensions from the plastics-enriched pellets and of synthetic plastics to discover the fluorescence profiles of these particles. A lambda scan is a spectral imaging mode that captures the full emission spectrum of a fluorescent sample across multiple contiguous wavelength channels rather than recording a single color channel. The scan records a 3D dataset (xyλ) where each image corresponds to emissions from a narrow wavelength band (e.g., 8.8 nm). Instead of filtering out neighboring light, the detector array records the complete intensity profile for every pixel across the chosen spectrum range, such that emission intensities of multiple particles within a large field-of-view across the whole spectra can be measured, analyzed and graphed. Computational post-capture analysis can be used to separate fluorophores with heavily overlapping emission spectra (like FITC and Rhodamine B) by mathematically decomposing the mixed signals using reference spectra fingerprints. Thus, fluorescence from other components in the brain can be minimized. Autofluorescence can be removed by identifying and subtracting intrinsic background autofluorescence to isolate the true target signal. Spectral unmixing starts with an operator-selected region (a particle in the suspension), propagates this region across all emission images in the dataset and generates an intensity-by-wavelength profile for each selected region (each particle). Thus, lambda scans can successfully be used to determine the unknown emission fingerprint properties of novel fluorescent objects, such as the putative fluorescence from plastics.

To determine whether particles in the plastics-enriched pellets (PEP) from human brain fluoresce, we performed lambda scans on suspensions of particles in the PEP from the two different dementia cases in which we had found fluorescent particles in the choroid plexus, whose histopathology demonstrated glossy perivascular deposits that fluoresced similarly to those in the choroid plexus, and whose PEP contained plastics by py-GC/MS (**Fig. 5**).

**Figure 5.**
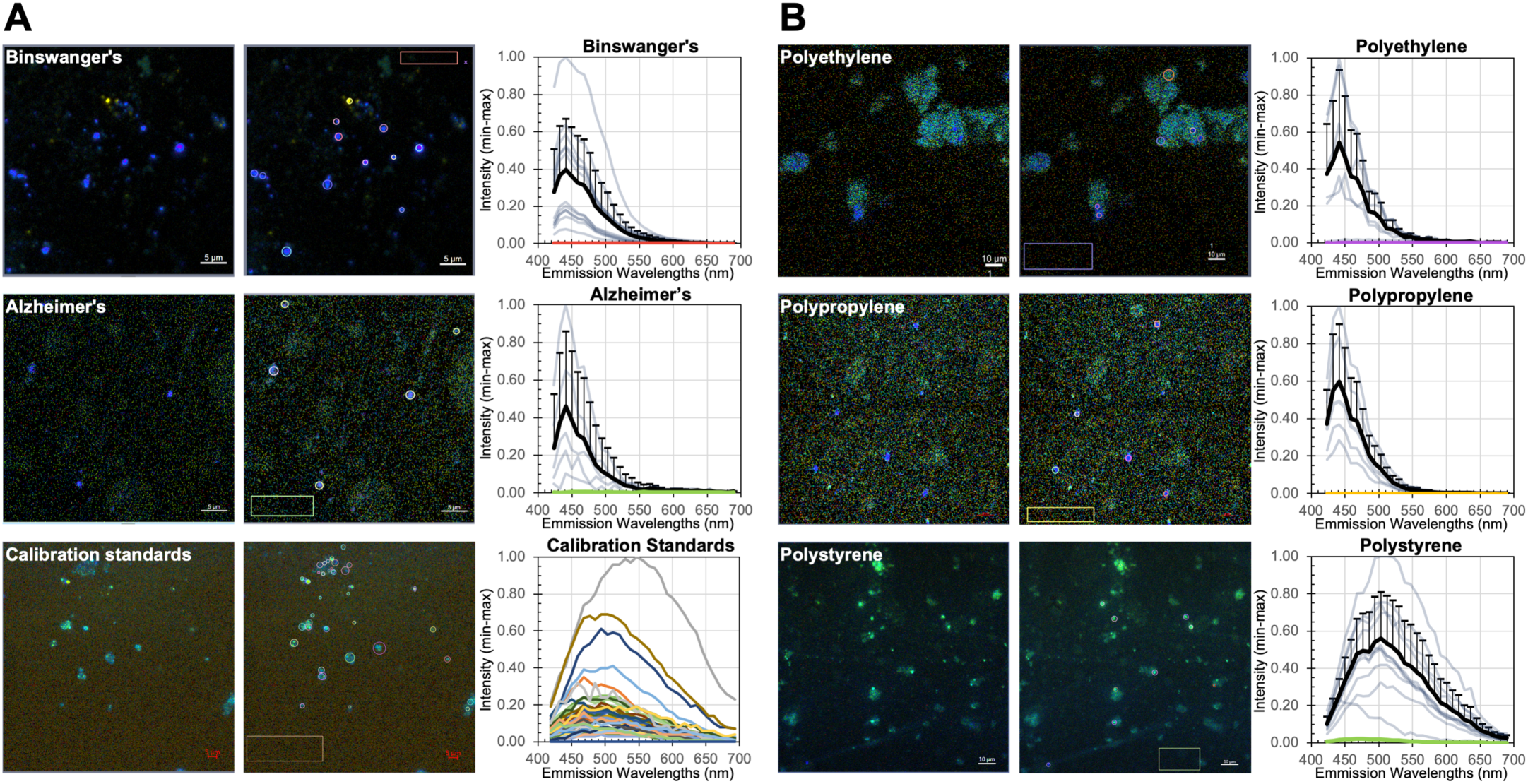
Hyperspectral analysis of particles and plastics. Fluorescence images and graphs of spectral profiles from aqueous suspensions of particles from **(A)** plastics-enriched pellets of brain tissue and from py-GC/MS calibration standards containing 12 different synthetic plastics; and from **(B)** synthetic plastics obtained from industry vendors. Suspensions mounted in coverslipped glass slides were excited by 405nm laser with emissions detected across the visible spectrum with the Zeiss LSM 980 confocal laser scanning microscope equipped with GAsP 32 channel detector running Zen Blue software. The 405nm laser was set at 4% power. Emissions were collected at 8.8nm intervals and the full 32-channel image stack superimposed. Left panels in each row show the image stack; middle panels, the same field of view with circle ROIs and background square regions indicated; right panel, graphs of intensity/wavelength across the 32 channels. ROIs were selected based on a visible particle in the stacked images, and sized based on the smallest circle to include signal and as little background as possible. The intensity values were normalized by Min/Max. **(A)** Top panels: a case of Binswanger’s disease, collected with pinhole at 1 A.U. (49µm) with a 63x oil immersion, 1.4 NA, Plan Apochromat objective with 3.1x zoom, giving a pixel size of 0.085µm^2^ with detector gain at 804V, 0 offset, Digital gain of 1. Mag bar = 5 µm. Graph shows ROI intensity values from individual particle (*blue-gray*), their averages (*black*, error bars = Std. Dev.), and the background measurement (*red)*. Middle panels: a case of Alzheimer’s disease with images collected with the same settings (n_particles_ = 5) as for the Binswanger’s case above. Mag bar = 5 µm. Bottom panels: Calibration standards used for py-GC/MS. Images collected with the same settings as above except no zoom, pixel sizes 0.132µm^2^ and image size 1024µm^2^ (n_particles_ = 45). All particles in the suspension fluoresced but display different profiles across the spectrum. Mag bar = 5 µm. **(B)** Fluorescence images for synthetic polyethylene (PE) (*top row*, n_particles_ = 6), polypropylene (PP) (*middle row*, n_particles_ = 5), and polystyrene (PS) (*bottom row*, n_particles_ = 10), with corresponding averaged spectral intensity profile graphs (*black*, error bars = Std.Dev.), ROI spectral profiles (*light blue*), and background (*color*). For PE and PP with randomly sized particles, the same settings as in A were used except with pixel size 0.263µm^2^ to accommodate the variations in particle sizes. For PS, with all particles sized to 50nm or smaller, pixel dimension was 0.129 µm^2^ with image size 1024x1024. The brightness of these unlabeled particles allowed a lower detector gain of 724V.

Initially we scanned suspensions of PEP from two different dementia cases with each available laser (405, 440, 488, 514, 561, 594 & 639 nm) and captured lambda scans across wavelengths longer than each laser line. This revealed that the best excitation was 405nm with emissions also in the 400nm range, which is not a profile commonly found among autofluorescence entities in brain sections. We found that the peak emission for particles from either specimen was about 450nm, dropping to 0 emission between 512-556nm, depending on the intensity at the peak (**Fig 5 A**, top two rows). The wide error bars indicate variability between particles within this range, although all exhibited peaks at the same emission wavelengths (441 to 450nm) and a slight shoulder between 468-477nm.

We also performed lambda scans on the 12 plastics in the calibration standard (**Fig 5A**, third row). This revealed complex profiles, with every particle detected by transmitted light fluorescing but with many different emission profiles and intensities across the spectra when excited by 405nm laser. Thus, we expect that all of the 10 different types of plastics found in the brain that were also in the calibration standard will be fluorescent and thus potentially be detectable in tissue under these imaging conditions. However, the best test will be of each of the different plastics in the calibration standard to be tested individually, which is beyond the scope of the current study. To start, we explored three, polyethylene, polypropylene and polystyrene all of which fluoresced with 405nm wavelength excitation.

To learn whether specific plastics may have unique hyperspectral profiles and whether these could be used to identify them in our specimens, we obtained single species preparations of synthetic plastics (polyethylene, polypropylene and polystyrene), made aqueous suspensions and performed lambda scans on them (**Fig. 5B**).

Polyethylene had large particles in the commercially available powder (**Fig. 5B**, top row). Intensity of ROIs from within the particles peaked between 441nm and 459, with two shoulders at 477 and 503, and fall off by 556 to 0. Polypropylene also peaked at 441nm, with a shoulder at 468nm and fall off at 565nm (**Fig. 5B**, middle row). Whether these minor differences are enough to distinguish between them in tissue is not clear. In contrast, polystyrene was distinctive from PE and PP (**Fig. 5B**, bottom row), with a peak at 512nm, an early shoulder at 485nm and a fall off to 0 not until 680nm. This profile could clearly be distinguished from PE and PP.

These results demonstrate that the synthetic polymer materials tested exhibit detectable fluorescence under 405 nm excitation, with distinct emission profiles, not predicted based on their theoretical chemical structure. This property will potentially allow them to be detected, and possibly typed, in human tissue.

### Positive and Negative controls: Polystyrene and water

Since all our plastics were resuspended in water, we tested this water for presence of plastics by negative stain EM and by confocal laser scanning microscopy (**Fig. 6**). To ensure that plastics would be detectable with our imaging parameters, we used 50nm polystyrene beads as a positive control. These are smaller than the plastic shards previously reported to be 40x200nm. Polystyrene beads suspended in water were easily imaged after negative stain on formvar-carbon coated glow-discharged 200 mesh copper grids by electron-microscopy (**Fig. 6A**). Similar grids processed in parallel with water only followed by negative stain displayed no particles (**Fig. 6B**). Un-labelled polystyrene beads mounted on grids in parallel but without negative stain and imaged by confocal had the typical fluorescence profile of PS bead suspensions on microscope slides (**Fig. 6C**), whereas water alone had a few particles detected by the PMT, although those that were seen lacked detectable fluorescent signal under similar imaging parameters. In addition to the controls shown here, we also tested regions adjacent to tissue in stained slides, and regions with no tissue within the section, few if any fluorescing particles were found on any slide. We also performed z-stacks on particle suspensions, and found all particles within the aqueous chamber containing the suspended particles and none above or below it. If there had been contaminating plastics in any of the staining solutions, we would have seen them, which we did not. We thus concluded that our imaging parameters were sufficient to detect nanoparticles, and that the water used for particle suspensions did not include any plastic particles and was not producing the intensity profiles, shown in **Fig. 5** above.

**Figure 6.**
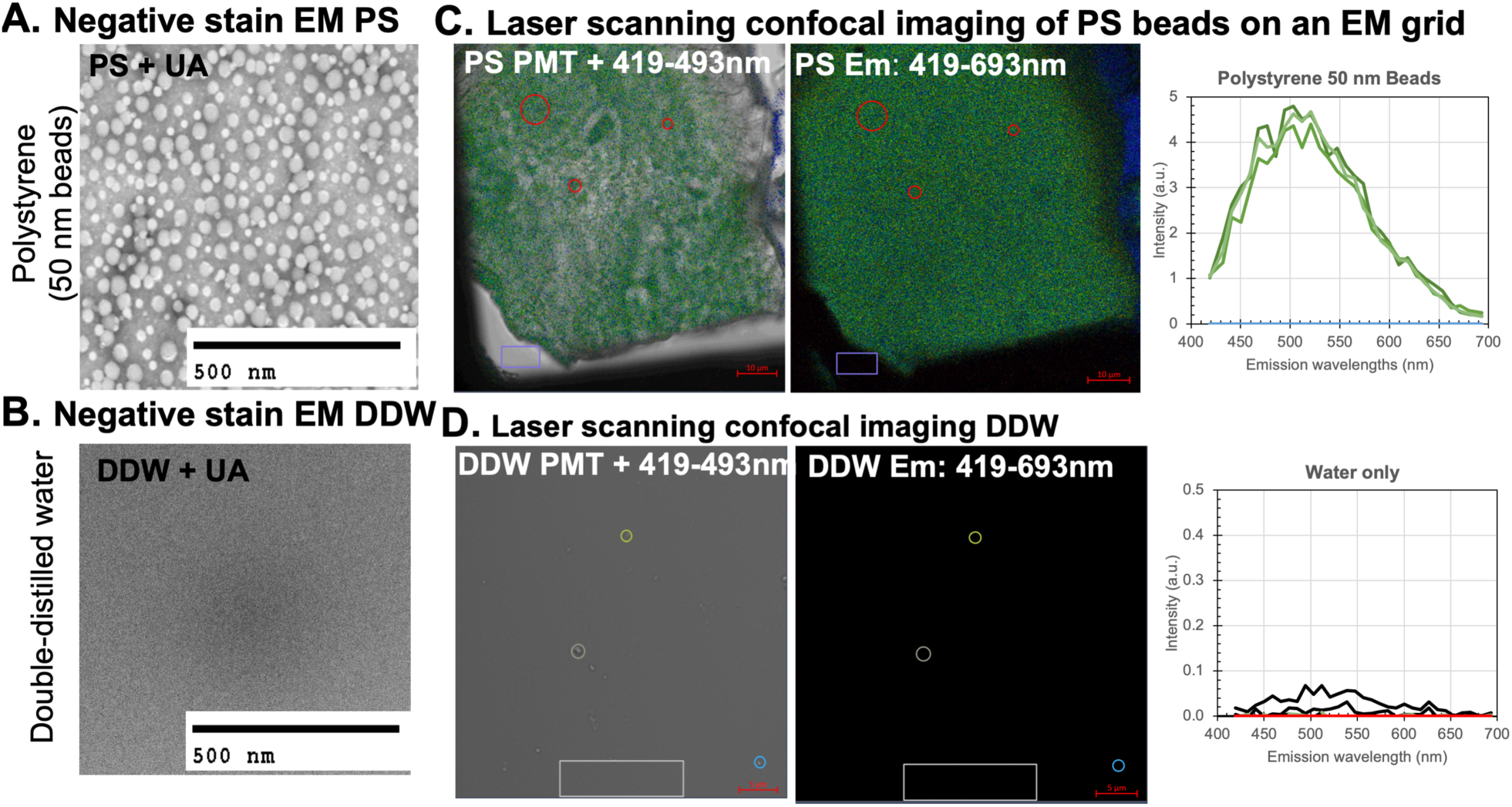
Positive and negative controls. (A) Negative-stain electron-microscopy detects 50nm and smaller beads in the synthetic polystyrene sample. Mag bar = 500nm (B) In contrast, no objects are found in carbon/formvar coated glow-discharged grids prepare in parallel. Mag bar = 500nm (C) Laser-scanning confocal imaging detects polystyrene beads on a grid not stained with uranyl acetate as particles in the gray scale PMT image superimposed on the 32-channel fluorescence (left panel). These particles emit in the 450-550 range (middle panel image of the 32 channels and right panel graph of emission profile). Vertical axis is in arbitrary units calculated by Zen. Mag bar = 10 µm. (D) While the PMT detects a few tiny particles in DDW, these are not fluorescent. Note the vertical scale of the graph to the right is from 0-0.5, a tenth smaller than that in C. Mag bar = 5 µm.

### Subcortical blood vessels display particles that fluoresce like plastics

Confocal imaging of blood vessels demonstrated multiple particles that fluoresce with a similar profile as plastics, and that this profile is unique and not shared by collagen, RBCs or tissue (**Fig. 7**). Lambda scans of this section, which includes perivascular collagen (green circles and yellow ‘c’), RBCs in the vessel lumen (red circle), adjacent brain tissue (pink circle), non-tissue regions (white circle) and tiny particles (blue circles and blue arrow) in the perivascular space that emit only in the mid-400s (blue **Fig. 7A**). Such images detected fluorescing particles and also showed tissue architecture. When the channel with emissions of 465-472nm is displayed as a single image, distinctive blue-fluorescing particles appear (blue arrow) (**Fig. 7B**). Spectral un-mixing shows that these particles have a peak fluorescence between 441 and 500nm (blue lines) with little background from collagen (green lines), RBCs (red line) or tissue (orange line) line (**Fig. 7C**). These tissue components emit at wavelengths higher than 512nm.

**Figure 7.**
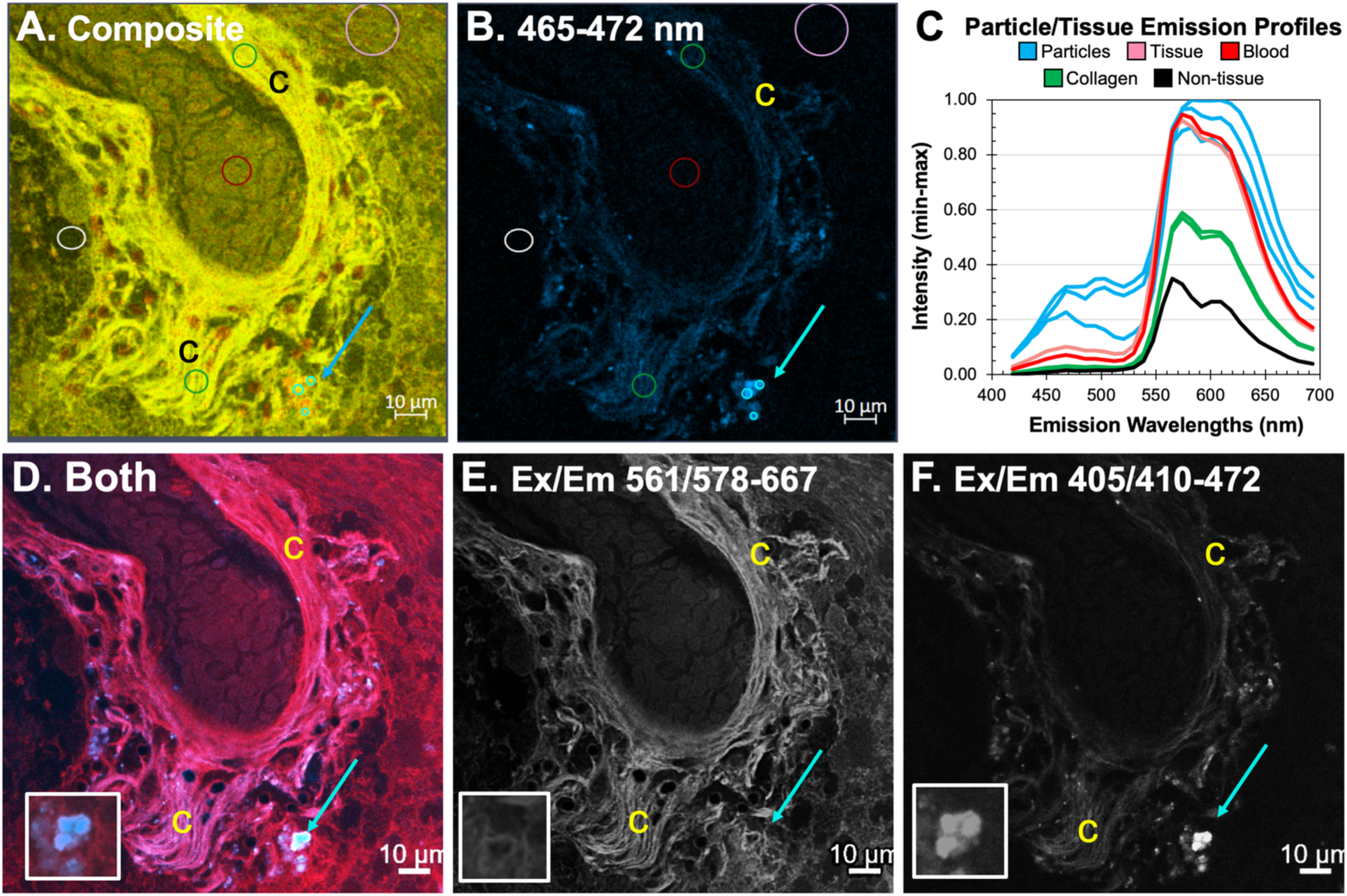
Application of Hyperspectral Imaging to Tissue Sections of Human Brain. This is the same section of a subcortical white matter arteriole shown in Fig. 2F in brightfield, rotated 90° clockwise and at higher magnification. The section was stained for hemosiderin with Prussian blue and counter stained with nuclear fast red. The deposits did not stain for iron. (**A-C**) Lambda scans of emissions in Zeiss LSM 980 across the visible spectrum, as for synthetic plastics shown in Figures 5-7. Excitation by 405 laser at 3.5% power, 20x 0.8 NA Plan-Apochromat objective with 3x zoom, pixel dimensions 0.276, A.U.at 1 (28µm), Detector gain at 797V and 16 averages. (**A**) shows the full 32-channel stack, and (**B**) shows only the image at emission of 465-472nm, the peak for PE and PP. Circles indicate ROIs captured for emission intensity profiles which show that under these conditions, particles in the vessel wall emit in the 400nm range, while tissue, and blood do not. Notably collagen (yellow letter c) in the vessel wall does not emit in this range, and also has a different shape—long thin fibrils that polarize rather than tiny round aggregated particles. Blue arrows point to a cluster of particles in the vessel wall that emit in the 450-500nm range with 405 excitation. Mag bars = 10 µm. **(D-F)** Two-color image of the same field in the same section demonstrating the traditional fluorescence microscopy with appropriate filters might be sufficient to detect plastic-like particle in human tissue. Images were captured with multidimensional imaging on the Zeiss LSM 980 by exciting first with the 405 laser and programming the detector to acquire emissions from 410-472nm; and capturing a second separate image, exciting with the 561nm laser and acquiring emissions from 578-667nm. (**A**) shows both images superimposed; (**B**) shows the single channel 578-667nm emission after 561 excitation; (**C**) shows the single channel 410-472nm emission after 405nm excitation. Insets in lower left of are arbitrary enlargements of particles indicated by blue arrows. Mag bars = 10 µm.

Applying these unique spectra to design a two-color image, we acquired an image of the same vessel but instead of a lambda scan with one 405 laser, we obtained images in two channels with two different laser excitations: 405 and 561nm (**Fig. 7 D-F**). We set the detector for each channel to collect the emissions excited either by the 405 laser for particles (410-472nm) in channel 1, or for the tissue by the 561nm laser (578-667nm). in channel 2. This schema collected a 2-color multidimensional image (**Fig. 7D**) with each emission according to the spectral profile of either tissue (**Fig. 7E**) or particles (**Fig. 7F**), as identified by lambda scans. Note the blue particles in 410-472 channel. While there are three large ones in the right lower region of connective tissue, there are many more smaller ones throughout the vessel walls. Also note that the collagen signal does not overly the particles fluorescent and give no signal to them.

### Glossy deposits fluoresce like synthetic plastics in histologic sections of brain

To determine if these fluorescent particles represented those initially observed as glossy yellowish deposits adjacent to blood vessels in the brain, we went back to the same slide with the section imaged in **Fig. 7** by laser confocal, to find the corresponding blood vessel and captured bright field (BF) image of it on a different microscope (**Fig. 8A-B**). This was necessary because the confocal microscope did not have a color camera for BF imaging, and thus confocal fluorescence images were captured without prior knowledge of their appearance in BF images. Correlations could only be done by capturing images of the same field in the same section with two different microscopes and aligning them post-capture (**Fig. 8**). This forced us into a type of “blind” experiment, where whether the glossy deposits fluoresced like plastics was not known *a priori*. Indeed, we found that the bright blue-fluorescing particles emitting under 405nm excitation at 465-472 also appeared as glossy yellowish deposits in BF as previously noted (**Fig. 1 and 2**) (**Fig 8 A & B,** boxed regions and enlargements in **Fig. 8C**)). Only one of the three fluorescing particles correlates closely with the yellowish deposit in the BF image, although there appears to be slight yellow discoloration under the adjacent nuclei stained with nuclear fast red that are invisible in the 465-472nm fluorescence channel. These red nuclei obscure the glossy deposits in the BF image although some yellowish discoloration can be seen behind them.

**Figure 8.**
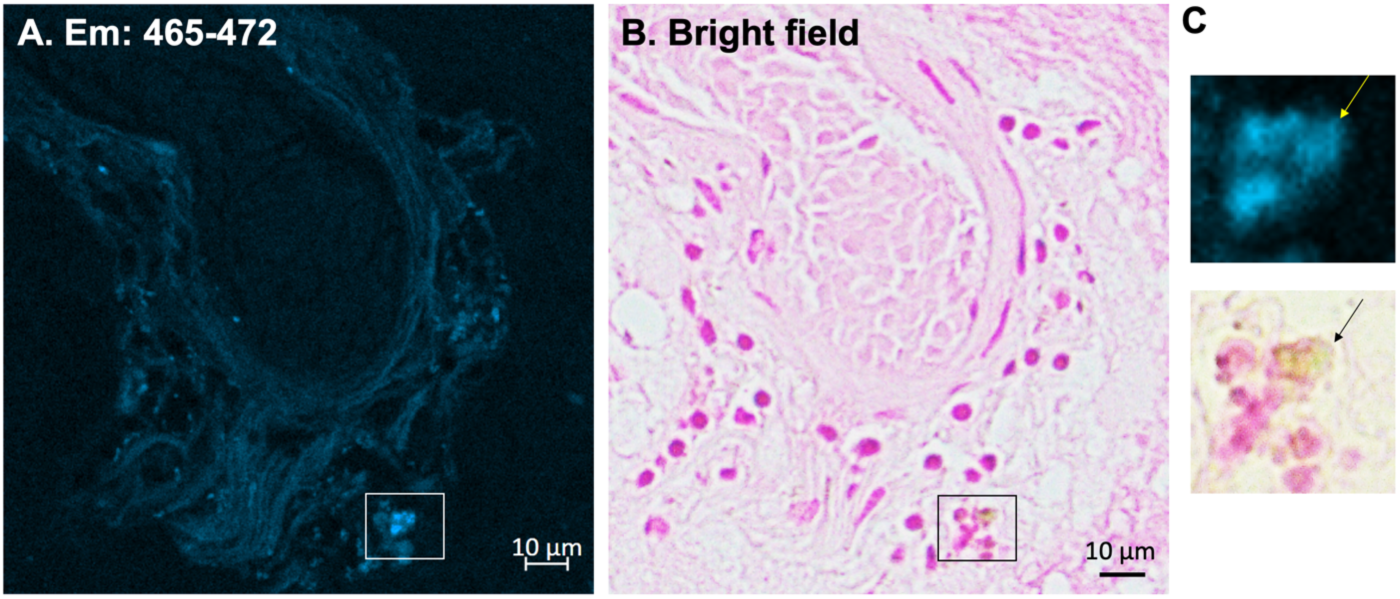
Particles with plastics-like spectra are coincident with yellowish deposits seen in IHC-stained sections. (**A**) The fluorescence emission at 465-472nm from the confocal laser lambda scan at 405nm excitation shown in Fig. 7 compared to **(B**) a bright field image of the same vessel in the same slide. The slide was stained with Prussian blue and nuclear fast red which shifts the collagen and formalin autofluorescence to longer wavelengths. The yellowish deposits are slightly less noticeable with this stain than with IHC shown in Fig. 1, while the fluorescence of particles is less easily detected with the IHC stain and its hemotoxylin counter, which is blue and gives a blue background. No blue ferrocyanide staining is apparent in this section, and blue ferrocyanide stain does not give any fluorescence in this wavelength. Arbitrary enlargements of the boxed regions in (A) and (B) show the particles distinctly in both imaging modalities

Because Prussian blue staining colors tissue blue, we also examined the hemosiderosis liver controls by lambda scans and found that the ferrocyanide, an iron stain, obscured fluorescence and did not induce a blue emission with 405nm laser excitation. We also performed lambda scans on non-tissue containing regions of the same slide and found minimal emission and very few particles. Finally we checked that the fluorescing particles were within the tissue by obtaining z-stacks. Particles were found within the section and not above or below it, demonstrating that they were not contaminants falling on top of the specimen during processing, or particles on the glass slide or coverslip.

## Discussion

In this study, we evaluated hyperspectral confocal fluorescence microscopy as a method for detecting and localizing micro- and nanoplastic particles within human brain histologic section. Synthetic polymer standards exhibited reproducible fluorescence under 405 nm excitation, and spectrally similar particles were observed in brain specimens in which polymers were detected by py-GC/MS. We used synthetic plastics as gold standards for both py-GC/MS and for confocal fluorescence microscopy. We found that all 12 plastics in the calibration standard fluoresced. This fluorescence was excited by 405 laser with variable emission profiles. Thus, different types of plastics may be identifiable by their fluorescence just as they are by their mass spectroscopy profiles, although much work will need to be done to define unique spectra for each type of plastic as fluorescence similarity alone may not establish polymer identity.

PS and PE had similar spectra and might not be distinguishable by this technology. The particles identified in tissue had spectra most similar to PE and PP and not to PS, which had a longer emission wavelength. Many experimental models aiming to understand effects of plastics on tissues in vitro or in animal models have relied on PS, partly as it is available from many sources^32,33^. Whether this plastic is representative of the plastics actually found in humans is a question. Designing experiments based on the real world data from human pathology will undoubtedly improve understandings.

There are many limitations in this study. For example fluorescence does not independently establish polymer identity. The field is rapidly advancing with new methodology to help clean up pellets further than those described here. Only two index cases were studied in depth here as our aim was method development not epidemiology. Thus no statistical information about prevalence is possible or appropriate at this time. It may be that the size, shape and surface charge of plastic particles is more significant for health impacts and biological responses than the specific plastic type. While some characterization techniques allow for quantification of the amount of specific types of plastics within tissue samples, fluorescence microscopy may be less specific but more sensitive to smaller fewer particles and informs on their tissue distribution. This new methodology represents a trade-off between the specificity and quantification of py-GC/MS versus the higher sensitivity and resolution of fluorescence imaging which proved information about localization of minute plastic particles within tissue architecture. In the brain, such precise localization may be more important for health than overall quantity measured in large chunks of brain matter.

The novelty of this area presents a limitation, as relatively little prior research is available to inform the study’s framework and interpretation. For example whether other non-biological particles may be present in brain that also fluoresce like plastics is possible but unexplored. What the source of the fluorescence may be in plastics that are composed of single bonded carbons—is fluorescence emanating from an additive, such as the commonly used but not covalently attached optical brightening agents, colorants and bluing agents? These would leach out over time and thus not be a continuing reliable signal for all plastics. If not coming from some specific chemistry within each plastic type, how stable is the profile, and will that be useful for identification across cases and conditions? And how wide is the variation of abundance, types, sizes and shapes of plastics in human brains? We don’t yet know how large a cohort would give a statistically useful sample size. It is likely that multiple different hydrophobic polymers aggregate in the aqueous tissue environment to produce these fluorescing particles. There is much work to be done.

A key unresolved question is whether the presence of plastics within the body has adverse health consequences or merely represents a benign bystander phenomenon. While the possibility of harmlessness may remain for some situations, there is thirty-year evidence from a “natural” experiment that micro-nanoplastics can have severe harmful effects that do not resolve over time. In 1996, a number of workers at a textile factory in Rhode Island that manufactured upholstery from microfibers were found to have an unexplained interstitial lung disease (ILD)^34^. Biopsies were performed and examined by pathologists which revealed a consistent and unusual non-granulomatous lymphocytic inflammation. Workers at another factory in Ontario, Canada, manufacturing the same upholstery, also came down with a similar type of ILD. Both factories used “flocking” of microfibers made of PE, PP and vinyl (a polyamide). The CDC ultimately named this ILD “flock worker’s lung.” Among the 165 members of the cohort of all the Rhode Island factory workers, seven (4%) were identified with “flock-worker’s lung.” The mean age for these seven was 41 years (range: 28-57 years); six were men. At the Rhode Island plant, the crude incidences of “flock-worker’s lung” and of all ILD were 10.5 cases per 1000 person-years and 15 cases per 1000 person-years, respectively. General population estimates for age- and sex-specific incidence of pulmonary fibrosis/idiopathic pulmonary fibrosis (PF/IPF) and for sex-specific incidence of all ILD were obtained from an ILD registry for Bernalillo County, New Mexico^35^; Using these estimates and weights based on the demographics of the study cohort, standardized incidence ratios for pulmonary fibrosis and for all ILD were calculated. Ninety-five percent confidence intervals (CIs) for these estimates were derived by exact Poisson calculations. The two clusters described in these reports together constitute the largest unexplained continuing outbreak of non-granulomatous chronic diffuse ILD in adults under investigation by CDC (https://www.cdc.gov/mmwr/preview/mmwrhtml/00049601.htm) . The cases in Ontario were followed up in 2013, and no one had recovered fully^36^. Respirable dust, characterized by both phase contrast microscopy and scanning electron microscopy with energy dispersive x-ray analysis, consisted of particles with physical structure and chemical composition similar to those of bulk samples of the finished material; a substantial number of respirable-size fragments of nylon plastic also were present. None of the affected people completely recovered. Hence we have known for thirty years that some types of plastics together with their finishing components in some situations do cause harm.

In conclusion, we demonstrate that synthetic polymer materials can exhibit detectable fluorescence under hyperspectral microscopy and that particles with similar spectral properties can be visualized within human brain histological sections containing polymers independently detected by py-GC/MS. This approach provides complementary information by preserving tissue architecture and enabling higher resolution to visualize micro- and nanoplastic particles in relation to surrounding cells, blood vessels, and pathologies. When used alongside established techniques for polymer identification, hyperspectral microscopy may provide a valuable tool for investigating the localization and distribution of potential micro- and nanoplastic burden within biological tissues. Further validation will be necessary to establish the specificity of these fluorescence signatures for definitive polymer identification. Because this technique is applicable to histopathologic sections, comparisons of the plastics’ presence, abundance, distribution, shape and possibly type with tissue damage at the cellular level is now possible.

## Acknowledgements

We are grateful to the Biological Imaging Facility in the Beckman Institute at California Institute of Technology in Pasadena, CA, for their superb confocal imaging equipment, and to Paul Webster of Oak Crest Institute in Monrovia, CA for thin-section preparation and electron-microscopy. We thank Louie Kerr and Carsten Wolff in the Central Microscopy Facility at Marine Biology Laboratories, Woods Hole, MA, for advice and assistance with negative stain EM, glow discharge on grids, and use of the JEOL CX200 electron-microscope. We are also indebted to the UNM College of Pharmacy for inspiration from Matthew J. Campen, and use of the BioAnalytical Chemistry Core of the NM-Inspires program and its Integrative Molecular Analysis Core with the Agilent 6890N GC-MS . We thank Dr. Rui Liu in this core for his advice, software access and training. We are thankful to the New Mexico Office of Medical Investigator for access to donated human brains for the UNM Brain Bank and to TriCore References Laboratories for expert embedding, sectioning and histochemical stains.

We have consulted several knowledgeable chemists whose advice advanced this study, including Ulf Berg, PhD, Professor of Chemistry at Lund University in Sweden, and Harry B. Gray, Professor of Chemistry at Caltech in Pasadena, CA, USA. We thank Lynn Rios, PhD, in the Bearer lab at UNM, Ying Wang, PhD, in the Biological Imaging Facility at Caltech for technical assistance, Joseph A. DeGiorgis, PhD, for help with the electron-microscope at MBL, and UNM medical student, Sean Pinon, for helpful discussions. Some of this work was presented previously as posters at the International Congress for Microplastics and Health in Santa Fe, January 2026, and at the annual national meeting of the American Association for Neuropathologists in Cambridge, MD in June 2026. Comments from experts in plastics and in neuropathology who reviewed this work during those presentations has been taken into account in this report.

This work was funded in part by National Institutes of Health P30AG08404 (Rosenberg/Bearer), P20AG068077 (Rosenberg/Bearer), F99NS139535 (Uselman); the Beckman Institute at Caltech (Bearer and Collazo); and the Harvey Family Endowment, UNM Foundation (Bearer).

